# Permutation-calibrated stability discovery under p >> n : A leak-controlled Machine Learning framework identifies candidate proteomics panels in antiseizure medications-related side effects

**DOI:** 10.64898/2026.02.24.707860

**Authors:** Saman Hosseini Ashtiani, Markus Karlander, Sarah Akel, Johan Zelano

**Affiliations:** Department of Clinical Neuroscience, Institute of Neuroscience and Physiology, Sahlgrenska Academy, University of Gothenburg, Gothenburg, Sweden; Wallenberg Center of Molecular and Translational Medicine, University of Gothenburg, Sweden; Clinic of Neurology, Rehabilitation and Specialized Care at Home, Södra Älvsborg Hospital, Borås, Sweden; Region Västra Götaland, Södra Älvsborg Hospital, Department of Research, Education and Innovation, Borås, Sweden; Department of Neurology, Sahlgrenska University Hospital, member of ERN Epicare, Gothenburg, Sweden

## Abstract

We investigated whether the plasma proteome distinguishes people with epilepsy who report central nervous system (CNS) side effects from antiseizure medications (ASMs) from those who do not. In 161 patients profiled using proximity extension assay-based proteomics Neurology and Inflammation panels (∼1,447 proteins), we applied an ensemble leak-controlled machine-learning (ML) workflow based on LASSO (linear) and random forest (RF) (non-linear) with repeated nested cross-validation and stability selection. We engineered nested machine-learning workflows that embed permutation-based Monte Carlo p-value estimation directly within model training, enabling statistically calibrated feature discovery under high-dimensional, noisy proteomic data. Discovery phases were explicitly optimized for association and feature robustness, not prediction. The RF yielded a 61-protein candidate panel, for which an “exploratory nested RF” model achieved strong internal discrimination of CNS side effects (AUROC ∼ 0.92, 95% CI ∼ 0.86–0.96). The LASSO yielded a three-protein candidate panel all of which overlapped with those of the RF (SMOC2, TANK and IMPG1). Because per-protein testing across all 1447 proteins produced false discovery rates (FDRs) close to 1, we performed post-hoc, data-driven routed per-protein inference restricted to the 61-protein panel, identifying 13 proteins with FDR <0.1. Network and pathway analyses on the 61-protein panel highlighted immune, autoimmune and vascular-inflammation pathways (e.g. cytokine networks, JAK-STAT, T-cell–mediated responses), suggesting that pre-existing immune and inflammatory may modulate vulnerability to ASM-related CNS side effects. Technically, our contribution presents a resampling-based stability statistics and FDR control despite *p* ≫ *n* and weak global discrimination. This framework is model-agnostic and directly portable to other low-sample, high-dimensional, noisy omics settings by replacing the base learner (e.g., LASSO/RF/boosting) while keeping the same leakage-safe resampling and permutation-calibrated stability machinery to prioritize robust biomarkers over optimistic predictive accuracy. By explicitly separating robust discovery from post-selection exploratory modeling, the workflow provides a reproducible template for generating candidate panels when standard whole-proteome multiple testing is underpowered.

**Author Summary:** We studied whether patterns in blood proteins can distinguish people with epilepsy who experience central nervous system side effects from anti-seizure medications from those who do not. Working with 161 patients and about 1,447 measured proteins, we faced a common challenge in modern biology: far more measurements than patients, substantial noise, and strong correlations among proteins. To address this, we built a reproducible analysis template that prioritizes reliable discovery over optimistic prediction. Our approach combines two complementary machine-learning models and repeatedly tests them on held-out patients to prevent information from “leaking” from the test data into training. We then use repeated re-sampling and label shuffling to estimate how often each protein is selected just by chance, which lets us compute calibrated p-values and false discovery rates for machine-learning feature selection. This makes the resulting candidate protein panel easier to interpret statistically and less sensitive to random fluctuations in small datasets. Using this framework, we identified a 61-protein candidate panel and a smaller overlapping set of three proteins highlighted by both models, and then performed targeted follow-up testing within the panel. Because the workflow is model-agnostic and leak-controlled, it can be reused in many other omics studies with limited sample sizes.

## Introduction

Antiseizure medications (ASMs) are widely used in epilepsy management, but various side effects are inevitable.(1) Such side effects comprise cognitive and psychiatric disturbances and systemic issues. (1-4) The prevalence is estimated to be from 30 to greater than 95% in persons with epilepsy. (2, 5, 6) Central Nervous System (CNS)-related side effects due to ASMs involve cognitive symptoms, especially affecting memory and tiredness.(7-10) Predicting side effects is very complicated due to variable and individual-specific vulnerabilities with the same clinical and demographic characteristics.(10, 11) Many patients are also unable to report side effects, because of comorbidities or communication differences. Biochemical monitoring of brain health through blood samples is emerging in several neurological fields, including dementia. We hypothesized that certain plasma proteins or combinations of proteins could be associated with patient-reported side effects in epilepsy. If confirmed, this could indicate a potential for blood tests monitoring ASM safety and personalized treatment.

We analyzed OLINK proteomics data from a regional biobank study of epilepsy with both univariate outcome ANCOVA differential expression tests and leak-controlled machine learning classifiers to identify proteins associate with patient-reported CNS side effects. Our main question was if it is possible to develop a classifier derived from the plasma proteome to discriminate the patients who report CNS-related side effects and those who don’t. We used linear and non-linear machine learning classifiers, which are more sophisticated in capturing multi-dimensional and non-linear relationships between covariates compared with statistical tests that are often used to pinpoint differentially expressed entities for inferential purposes.

## Results

The demographic description of the cohort within each of the categories of CNS side effects, age and gender are presented in Table 1.

**Table 1:**
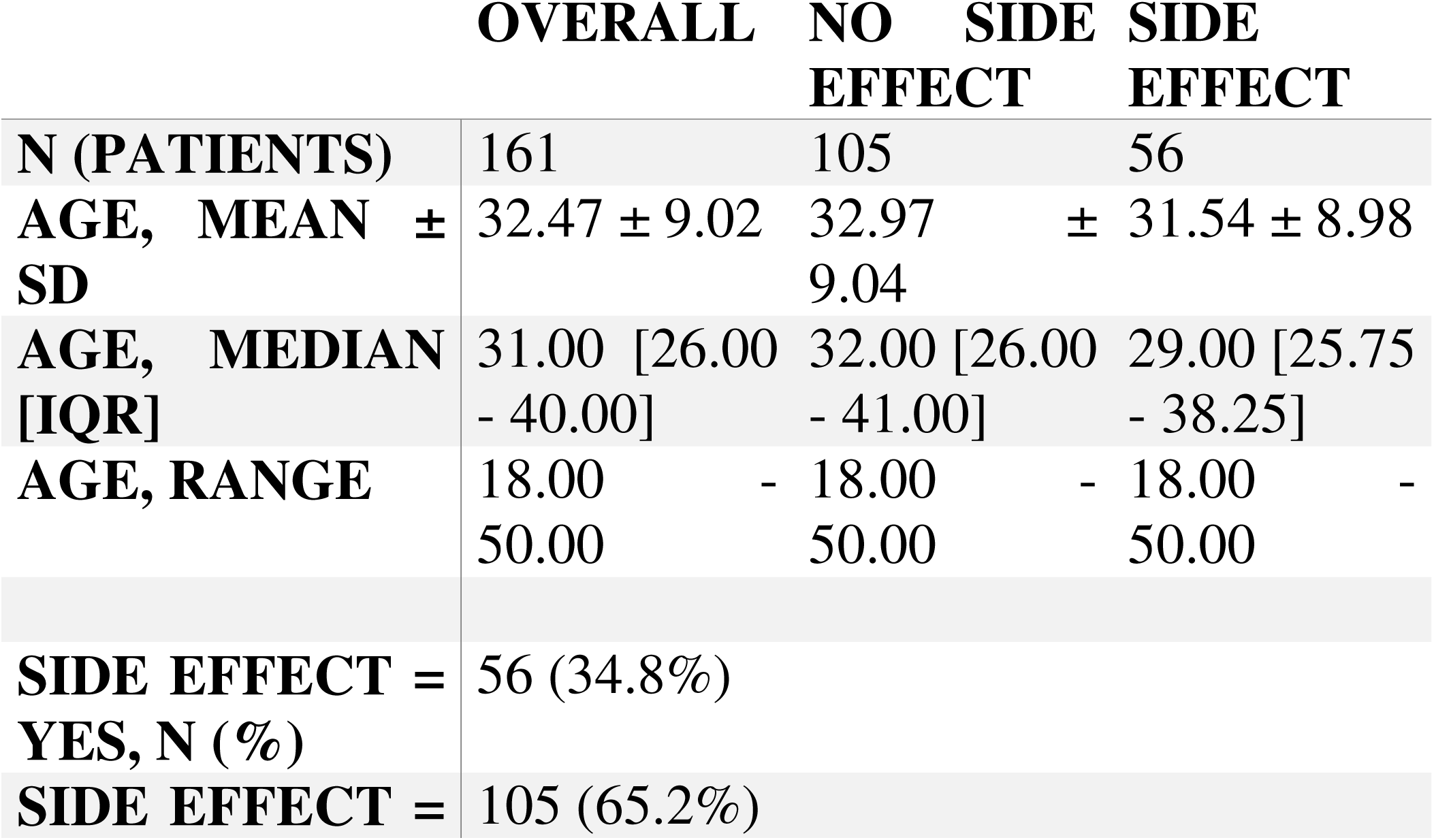
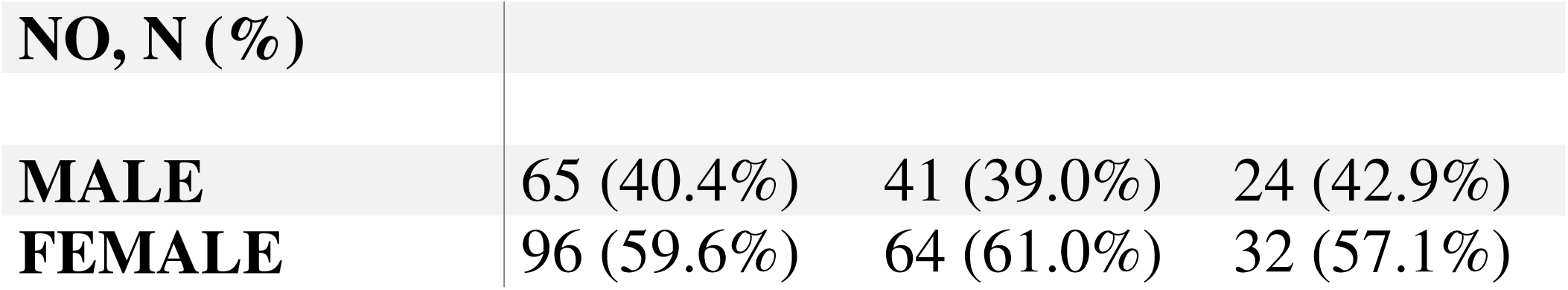
The cohort demographic table and CNS side effects groups with respect to age and gender.

### LASSO results

For the LASSO workflow, the regularization path shows the coefficient trajectories of the selected proteins IMPG1, SMOC2 and TANK with respect to the penalty increase (Fig 1A).

**Fig 1.**
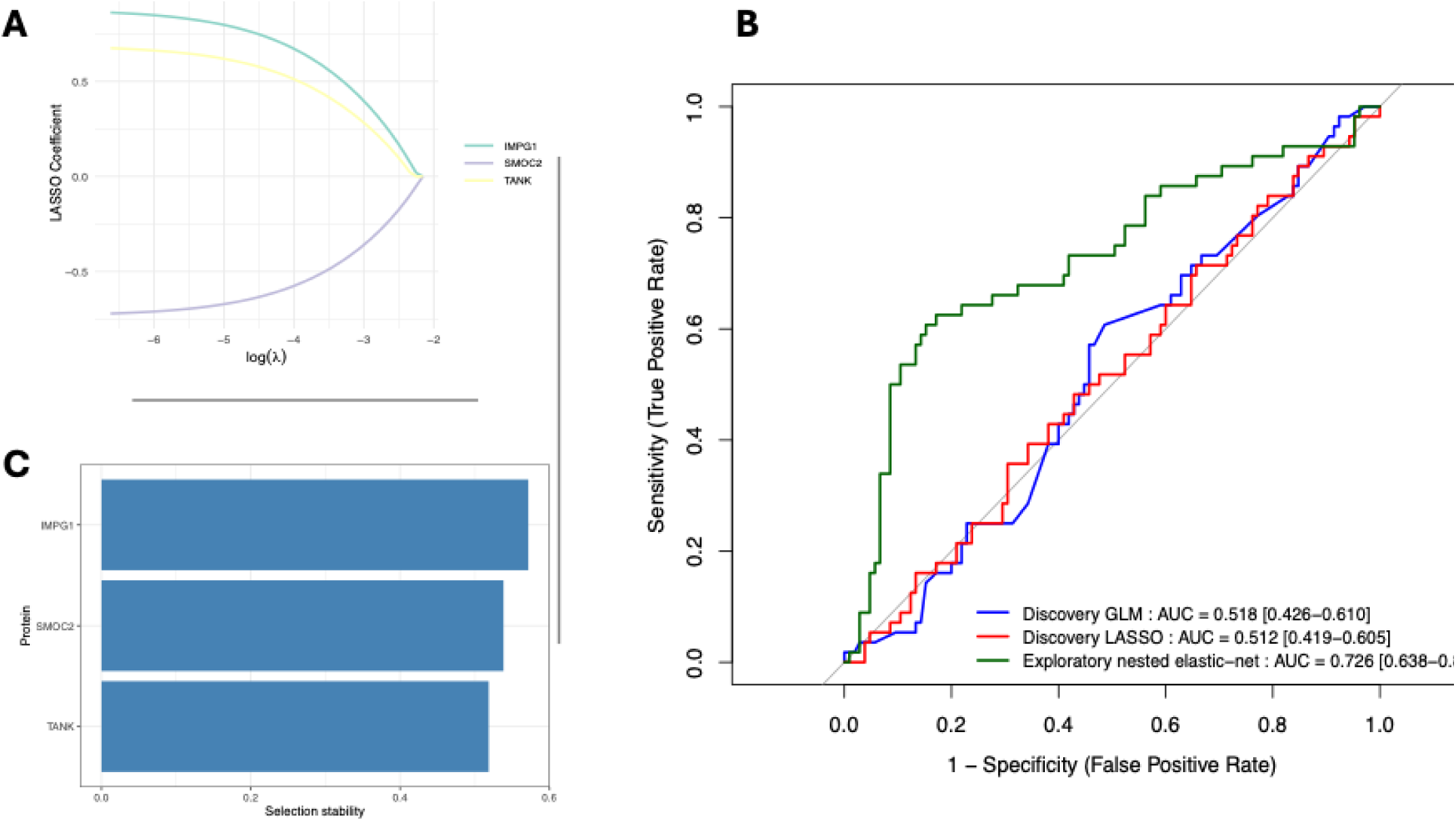
Discovery phase training and validation visualization for the LASSO workflow. A) LASSO regularization paths for selected proteins. LASSO regularisation paths for the three discovery-phase candidate proteins when fitting a penalised logistic regression model to all patients using only the candidate panel. Each coloured curve shows the estimated regression coefficient for one protein as a function of log(lambda), where lambda is the L1 penalty parameter. As lambda decreases (moving rightwards), coefficients move away from zero and enter the model at different penalty levels. The figure illustrates how the three proteins contribute to the penalised model across the regularisation path and complements the stability-selection analysis by showing the behaviour of coefficients in a single full-data fit. B) Overlay of ROC curves: Discovery and Exploratory comparisons. Comparison of ROC curves for three models evaluated using out-of-fold OOF predictions. The blue curve corresponds to a standard logistic regression model fitted to all 1447 proteins using the 10×10 repeated CV structure (“Discovery GLM”). The red curve corresponds to the discovery LASSO model fitted to all proteins with inner 5-fold CV for lambda and class-balanced weights (“Discovery LASSO”). The green curve shows the nested 10×5 elastic-net logistic model fitted only to the three discovery-phase candidate proteins (“exploratory nested elastic-net”). All curves are based on OOF probabilities. AUROC values and 95% confidence intervals (DeLong) are reported within each legend. The exploratory elastic-net AUROC is post-selection and should be interpreted as an internal exploratory validation summary rather than an unbiased estimate of external generalisation. C) LASSO selection stability of Discovery-phase candidate biomarkers. Selection stability of the three discovery-phase candidate proteins identified by LASSO stability selection. Bars show the stability S_j_ for each protein, defined as the proportion of 3000 resampled LASSO models (100 CV splits × 30 bootstraps) in which the protein was assigned a non-zero coefficient. All three proteins satisfy the dual criterion of high stability (≥ 0.5) and permutation-based FDR ≤ 0.20. Given the Monte Carlo standard error of stabilities (∼0.01), the observed values reflect genuine robustness of selection rather than resampling noise.

In an overlay of three ROCs to compare discovery and exploratory performances, discovery GLM and discovery LASSO models, both using all proteins, showed only modest discrimination, with AUROCs close to chance and overlapping DeLong 95% confidence intervals. The discovery GLM and discovery LASSO curves were broadly similar, indicating that shrinking coefficients and performing stability-based variable selection did not translate into a substantial gain in OOF performance when using the full 1,447-protein panel. The exploratory nested elastic-net model fitted on the three discovery-phase candidate proteins achieved an AUROC of 0.72 [0.64-0.81]. Consistent with its post-selection nature, this exploratory AUROC is best interpreted as an internal validation rather than evidence for robust external generalization (Fig 1B).

In the discovery-phase LASSO analysis, three proteins met our stringent stability and FDR criteria and were retained as candidate biomarkers (Fig 1C). Each protein achieved a selection stability ≥ 0.50 across 3,000 resampled LASSO models (100 CV splits × 30 bootstraps), while simultaneously controlling permutation-based FDR at ≤ 0.20. Given the Monte Carlo standard error of the stability estimates (∼0.01), these high stability values exceed what would be expected from resampling noise alone, indicating that all three proteins are consistently and reproducibly favored by the LASSO engine as discriminative features.

S1 Table shows proteins ranked by selection stability in the discovery-phase LASSO stability selection analysis. LASSO stability-selection statistics for all 1447 Olink proteins. For each protein, we report the selection stability S_j (proportion of all 3000 resampled LASSO models in which the protein had a non-zero coefficient), the corresponding mean selection frequency under label permutations (“NullFreq”), the one-sided Monte Carlo p-value p_j obtained from B_perm_ = 30 null permutations, and the Benjamini–Hochberg adjusted false discovery rate (FDR). Proteins with S_j_ ≥ 0.5 and FDR ≤ 0.20 (N = 3) were considered discovery-phase candidate biomarkers associated with CNS side effects and were used to define the reduced panel for the exploratory elastic-net analysis. The stability metric S_j_quantified how frequently each protein was selected across 3000 resampled models; the Monte Carlo standard error of S_j_was on the order of 0.01, indicating that decisions based on a stability threshold of 0.5 were not artefacts of finite resampling. By comparing S_j_ to a label-permutation null and controlling the FDR at 0.20, we identified three proteins with both high selection stability and low permutation-based FDR, supporting their interpretation as candidate biomarkers associated with CNS side effects.

Importantly, leak-controlled pre-discovery AUROC estimates for both un-penalized logistic regression and LASSO models fitted to all 1447 proteins were close to 0.5, and the observed split-level AUROC distribution overlapped closely with its permutation-based null. This illustrates a key distinction between statistical association and predictive performance, i.e., the presence of robust, FDR-controlled associations at the protein level does not guarantee high individual-level discrimination. The limited sample size, high dimensionality, and collinearity among proteins constrain the attainable AUROC, even when some biologically relevant proteins are present.

The second-stage nested elastic-net analysis restricted to the three candidate proteins was explicitly internal validation and exploratory. Its AUROC provides an internal sense of how these markers might behave in a more parsimonious predictive model but does not constitute an unbiased estimate of generalisation performance, since the panel was derived from the full dataset.

### Random Forest results

The discovery-phase RF trained on all 1447 proteins showed near-random discrimination in pooled out-of-fold predictions (OOF AUROC ≈), consistent with the mean split-level AUROC distribution being centered near 0.5. Across the 10×10 repeated CV, RF permutation importance yielded a distribution of selection stability values, S_j_, with a subset of proteins showing high stability (S ≥ 0.5) and FDR < 0.10. This indicates that the global RF model operates in a noise-dominated data set, and it explains why FDR values remain high, even when empirical p-values from the symmetric null appear non-trivial. Under such conditions, increasing the number of permutations cannot realistically yield FDR < 0.10 this is a property of the weak underlying signal rather than of the permutation scheme itself. Despite the low global discriminative performance, the stability selection analysis revealed a small subset of proteins that were repeatedly selected as highly important across outer splits. Among the 1,447 proteins, 61 proteins exhibited stability S_j_ ≥ 0.5 demonstrating they were robustly ranked among the top 20 % most important features in at least half of the 100 discovery splits. S2 Table summarises, for each protein, its selection stability, selection count, empirical p-value, and FDR estimates. We therefore defined an RF candidate panel based on stability ≥ 0.5 and FDR < 0.1, yielding 61 proteins (Fig 2A), which we then evaluated in an exploratory 10×10 nested RF CV. The exploratory RF panel achieved an AUROC of 0.92 (CI [0.86, 0.96]). This nested cross-validation ensures that hyperparameter tuning does not contaminate performance estimation. These ROC overlays of discovery and exploratory RF demonstrate an internal validation and comparison between the corresponding ROCs (Fig 2B).

**Fig 2.**
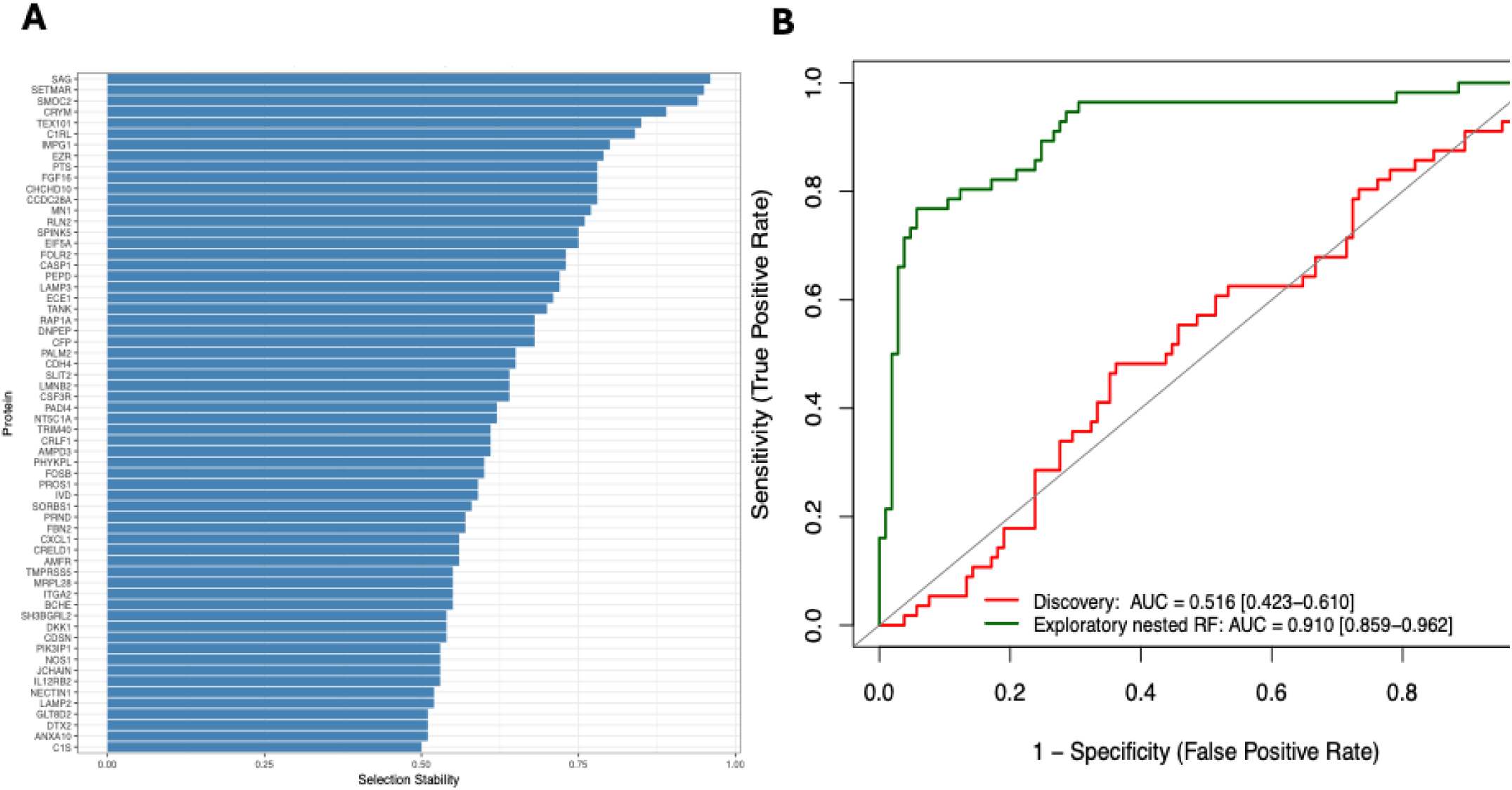
Discovery phase training and validation visualization for the RF workflow. A) Stable candidate biomarkers identified by Random Forest. Bar plot showing the stability S_j_ (proportion of discovery splits in which the protein appears in the top 20 % of RF permutation importances) for the stability-selected RF proteins (S_j_ ≥ 0.5) and FDR < 0.1. Bars are ordered by decreasing stability. This plot illustrates that a small subset of proteins is repeatedly selected as highly important across 10×10 outer splits, despite the near-null performance of the full-feature RF model. B) ROC curves for discovery versus exploratory RF models. ROC curves comparing 1) the leak-controlled discovery random forest trained on the full 1,447-protein panel (red) and 2) the exploratory random forest trained on the stability-selected protein panel (green). The discovery curve is based on pooled subject-level out-of-fold (OOF) predicted probabilities, obtained by aggregating predictions across the 10×10 repeated stratified CV and averaging repeated test-set predictions per subject. The exploratory curve is based on pooled subject-level OOF probabilities from a 10×10 CV procedure with inner-loop hyperparameter tuning on the fixed panel. The discovery RF shows near-random discrimination, whereas the exploratory RF achieves a high internal AUROC of 0.92 (DeLong 95% CI 0.86–0.96). The exploratory ROC is post-selection because the feature panel was defined using the full dataset discovery phase; therefore, the exploratory AUROC should be interpreted as internal, hypothesis-generating validation rather than an unbiased estimate of external generalisation.

Hierarchical clustering of the 61 overall candidate proteins revealed a proteomic structure across the 161 patients (Fig 3). Out of the seven protein clusters obtained using Euclidean distance and complete linkage, one of them showed heterogeneity in expression patterns. Notably, one compact cluster of 23 patients was characterized by uniformly low expression of CHCHD10, PALM2, NT5C1A, AMPD3, PRND, DNPEP, and LMNB2. Using all our available patients’ data we assessed if the 23 patients existed in any comorbidity or clinical sub-group, and they were not. We also checked the Olink plate sample sets to void the possibility of batch effect-induced clustering. None of the mentioned proteins were among those proteins with more than 30% of their samples being below Limit Of Detection (LOD) reported by Olink analysis experimentalists. We also made sure that all the proteins have passed the reported Quality Control.

**Fig 3.**
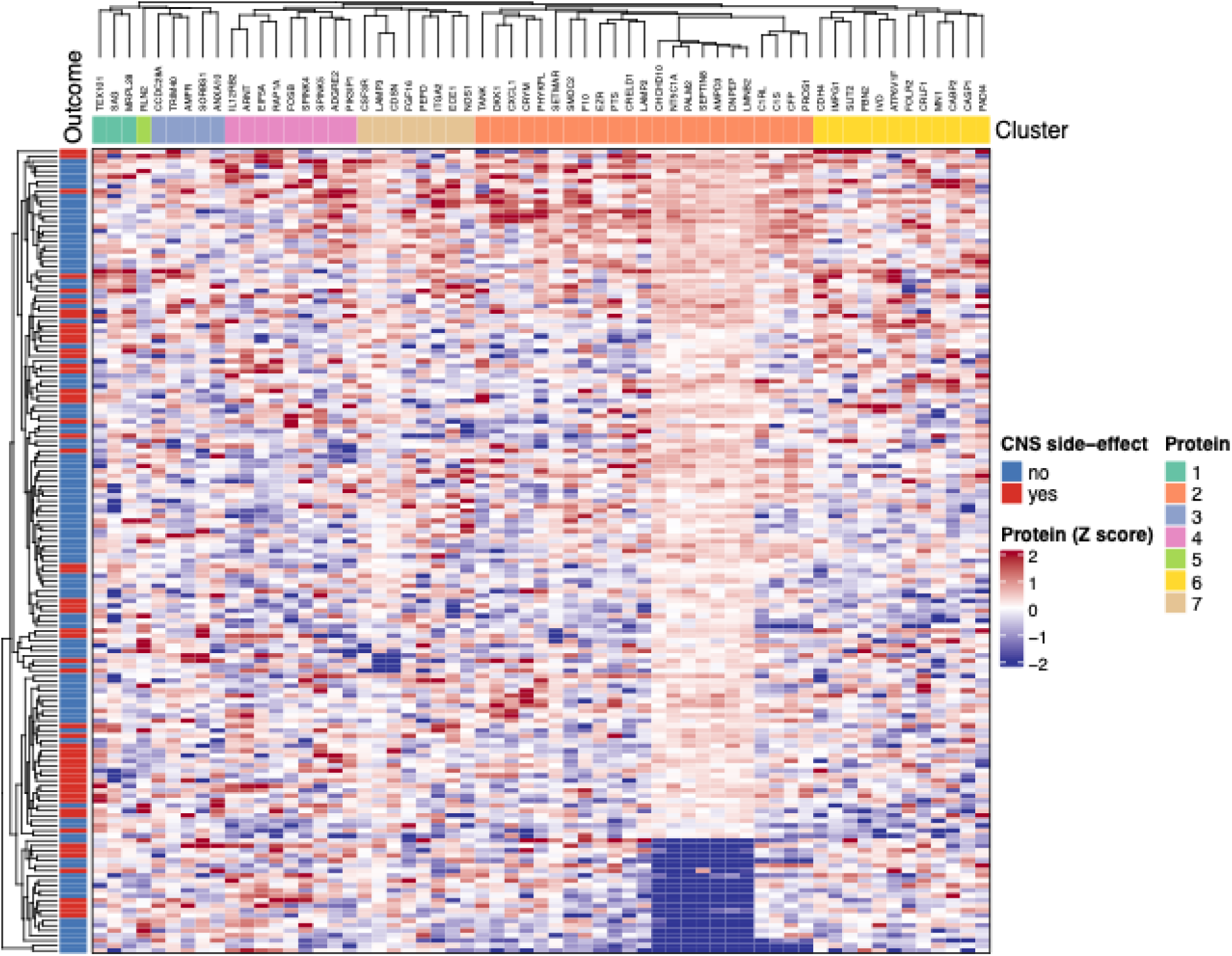
Hierarchical clustering of 61 RF stability-selected proteins that also contain the three LASSO stability- and FDR-selected proteins. Each column represents one protein and each row one patient (n = 161). Proteins were clustered using Euclidean distance and the complete linkage method, resulting in seven color-coded protein clusters (top bar). Row labels denote patients, colored by CNS side-effect status (“yes” = red, “no” = blue). Heatmap colors indicate z-scored protein expression (red = high, blue = low). A distinct low-expression protein cluster comprising CHCHD10, PALM2, NT5C1A, AMPD3, PRND, DNPEP, and LMNB2 is enriched for patients who were in patient cluster with side effects majority.

### Per protein differential expression analysis

We considered FDR < 0.10 as statistically significant, denoting moderate evidence after multiple testing adjustment. Under this threshold, 13 proteins (PHYKPL, IMPG1, SMOC2, RLN2, TANK, NOS1, PEPD, FGF16, TRIM40, CRLF1, AMFR, SAG, and CCDC28A) were identified as differentially expressed, three of which overlapped with the ML-workflow (SMOC2, TANK, and IMPG1).

Supplementary S3 Table shows that, outside the 13 FDR-significant markers, most proteins exhibit small standardized effect sizes and non-significant FDRs, consistent with a weak overall signal in this cohort. Nonetheless, the table provides a transparent view of the full post-hoc panel, enabling secondary analyses and cross-study comparisons even for subthreshold candidates.

The volcano plot in Fig 4 reveals that most candidate proteins exhibit modest covariate-adjusted log□ fold-changes and nonsignificant FDRs, in line with the weak global signal. However, 13 proteins occupy the “significant” region defined by FDR < 0.10, all with modest absolute effect sizes.

**Fig 4.**
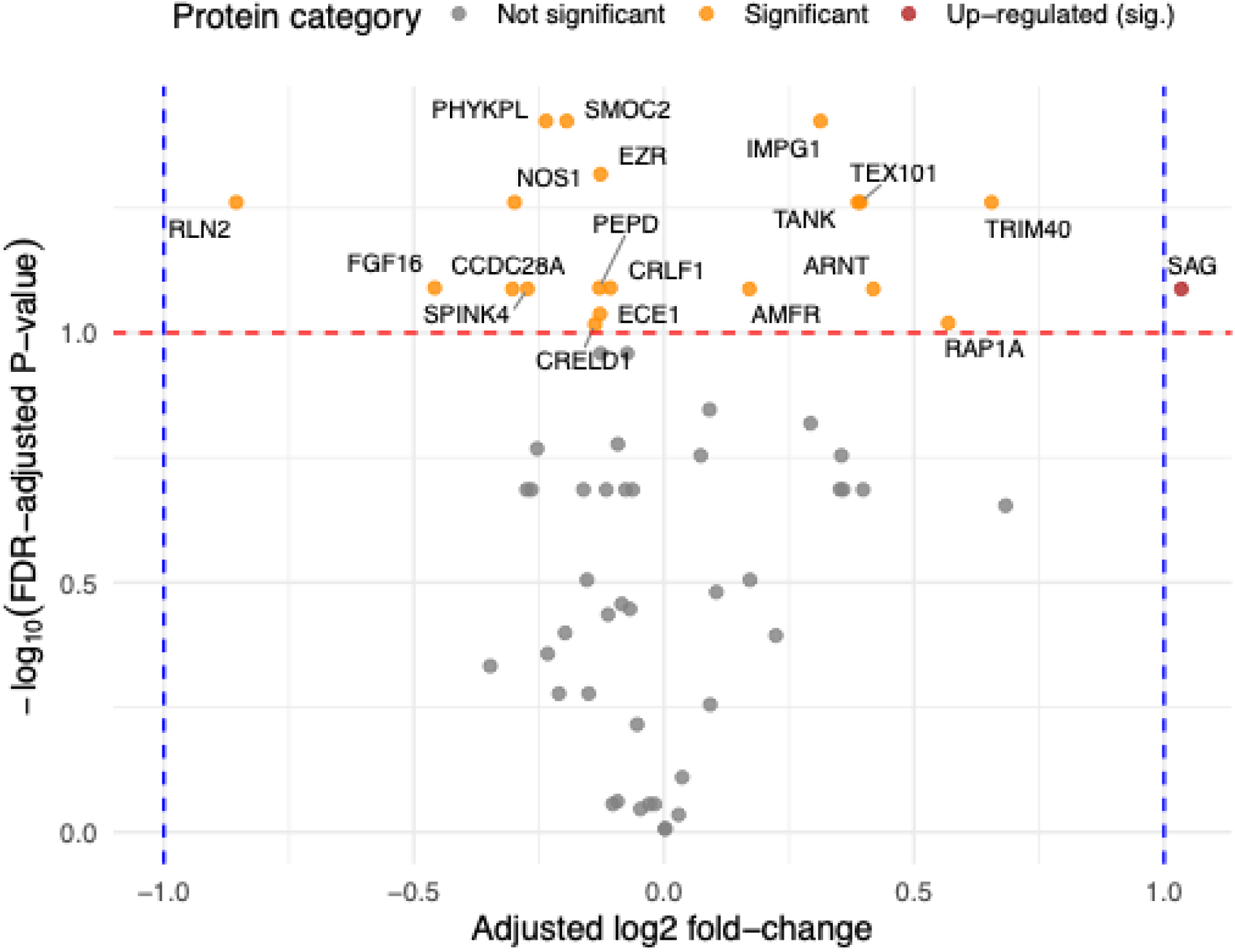
Volcano plot of covariate-adjusted log□ fold-changes versus FDR-adjusted p-values. Each point corresponds to one of the 61 candidate proteins, with the x-axis showing adjusted log□ fold-change between the “yes” and “no” groups (conditional on age and gender), and the y-axis showing –log□□ (BH-adjusted p-value). Vertical dashed lines indicate log□ fold-change thresholds, and the horizontal dashed line marks FDR = 0.10. Proteins are coloed by significance category (FDR < 0.10), and labels highlight FDR-significant markers, including SMOC2, TANK, and IMPG1, which also emerged as stable features in both ML workflows.

### Network analysis

The union of proteins identified by the three workflows (61 proteins with three overlapping ones) were used as the initial seed using STRING database online tools to achieve a PPI subnetwork. PPI evidence for the candidate panel was extracted from STRING as an edge list, where each row represents a putative interaction between two proteins (node1–node2) mapped to STRING protein identifiers (node1_string_id, node2_string_id). For each edge, STRING provided evidence-channel scores capturing distinct sources of support (genomic context, i.e., neighborhood on chromosome, gene fusion, phylogenetic co-occurrence; sequence similarity: homology; functional association: co-expression; experimental evidence, i.e., experimentally determined interaction; curated knowledge, i.e., database annotated and literature-derived evidence from automated text mining), along with an integrated combined_score summarizing overall confidence. This table was used to quantify interaction support across the candidate panel and to construct the PPI network underlying downstream network summaries and pathway interpretation (S4 Table). Using Cytoscape all the network connected components comprising 54 nodes were selected for the next step. Using Gephi modularity detection, seven modules were identified S5 Table. All the PPI connected components are visualized in Fig 5. We identified Module 2 (blue) as a distinct functional cluster enriched with inflammatory and immune-related protein candidates.

**Fig 5.**
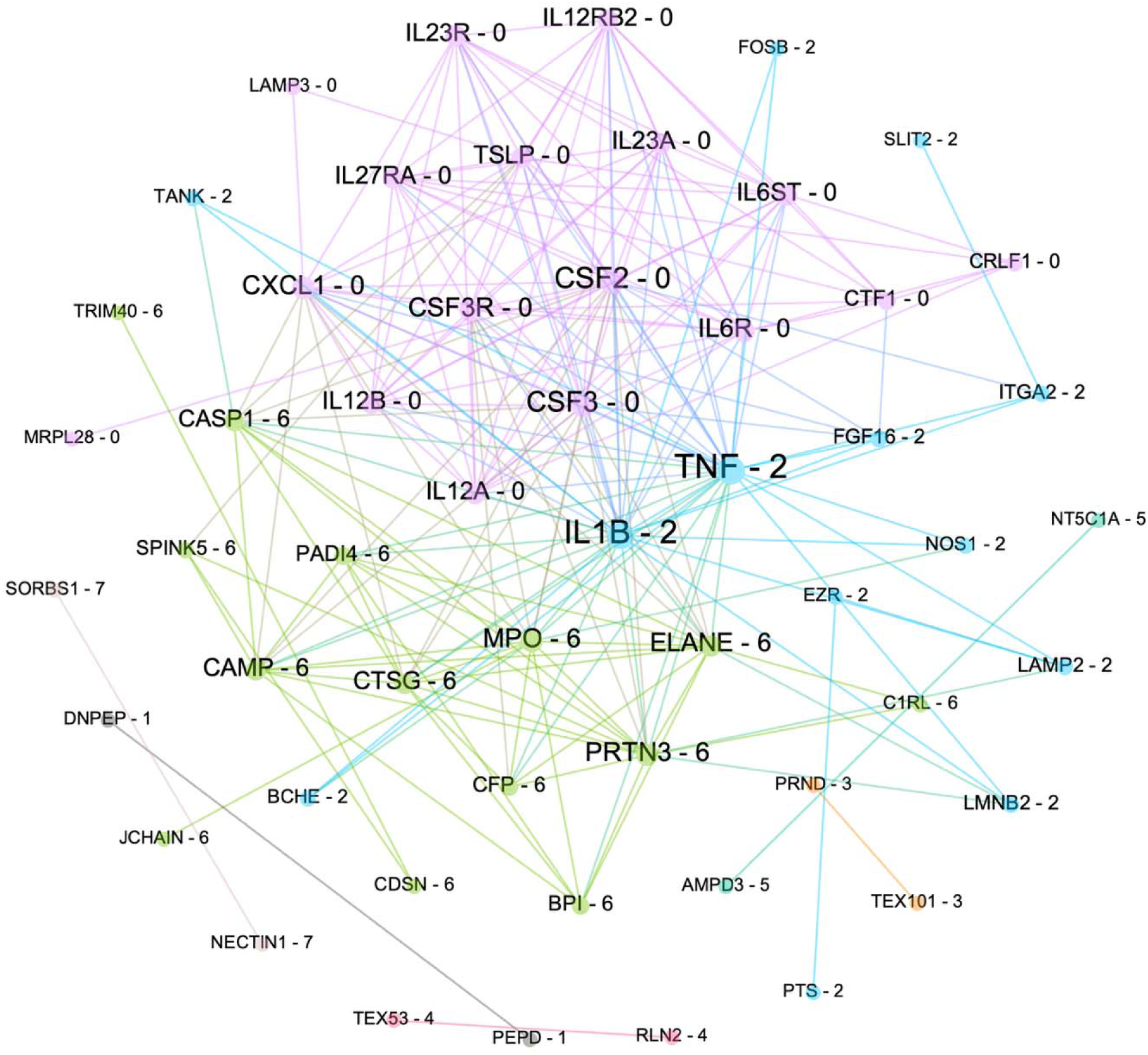
All components of PPI network analysis. Both large and small components are visualized. Each protein is tagged with a number indicating their module. The modules containing greater than two nodes, namely, module-0 (pink), module-2 (blue) and module-6 (green) underwent further literature review. The nodes in module are mainly involved in immunological and inflammatory pathways.

The results of enrichment analyses including Disease-Gene associations, Wiki-Pathways, KEGG-Pathways and Gene Ontology (GO) on all the proteins in the network connected components (#nodes=54) are presented in S1 Fig, indicating immune, autoimmune and vascular inflammation pathways and GO terms are mainly enriched with the candidate panel proteins.

## Discussions

### Computational

From a technical and ML perspective, this study contributes a generalizable strategy for biomarker discovery in the p ≫ n, noisy regime where classical per-feature testing across the full proteome is often underpowered and yields FDR near 1. We implemented a leakage-controlled discovery framework that treats model-based feature selection itself as a resampling-based statistic, i.e., selection stability is estimated under repeated CV with internal resampling, and its significance is calibrated against symmetric label-permutation nulls to yield Monte Carlo p-values and FDR control at the feature level. This “stability + permutation calibration” layer is model-agnostic and can be reused with other learners (e.g., boosting, SVMs, sparse PLS, neural nets with appropriate stability definitions), making it broadly applicable to small cohorts with thousands of correlated biomarkers. Finally, by separating a strictly leak-controlled discovery phase (optimized for robustness of association) from a clearly labeled post-selection exploratory phase (optimized for internal discrimination on the reduced panel), the workflow supports transparent independent validation studies. This data-driven study employed a dual leak-controlled ML workflow followed by post-hoc per-protein tests. A key novelty of our work lies in the engineering of the RF and LASSO workflows to compute permutation-based Monte Carlo p-values inside the cross-validation workflow rather than as a separate post-hoc layer. In the discovery phases of both ML workflows, we focused on association and robustness, not prediction. LASSO was chosen as the linear classifier engine because we had many proteins and relatively few patients, where variable selection and shrinkage were essential. In this scenario, L1-penalty regularizes the model to control variance and overfitting and simultaneously performs feature selection by shrinking many coefficients to zero(12). Overall, the workflow is deliberately conservative for discovery while prioritizing robustness and FDR control at the feature level and clearly separating exploratory predictive modelling from claims about out-of-sample clinical utility. As demonstrated by Varga et al., statistical association and predictive performance are fundamentally distinct concepts.(13) Bzdok et al., have technically discussed why inference and prediction need to be discriminated in biomedicine by showing that statistically important variables for inference are not guaranteed to produce high predictive performance. (14) Also, a recent machine learning work in rheumatoid arthritis also emphasized that biomarkers may show strong statistical associations with clinical outcomes yet exhibit poor predictive validity, reinforcing the notion that association is not prediction. (15)

Besides LASSO as the chosen linear classifier, to capture the non-linear interplays between the proteins involved in each group we opted for RF, because the bagging-based methodology of RF ensures stable and biologically meaningful feature selection, particularly in high-dimensional proteomic datasets(16).

Our post-hoc findings through per-protein analysis highlight the value of a two-stage, data-driven workflow in a setting where the global signal is weak and high dimensional. When we applied the same per-protein framework to all 1,447 proteins, FDRs were close to 1, indicating that naive univariate outcome ANCOVA (multiple linear regression) screening on the full panel is not informative in this dataset. We therefore performed the per-protein analysis as a post-hoc step restricted to 61 proteins identified by ML-based stability selection. Technically, this strategy has several strengths. First, feature pre-selection was based on nested cross-validated ML stability, not on univariate outcome p-values, minimizing leakage between the ML and per-protein stages. Second, the post-hoc inference was heterogeneity-aware, i.e., each protein was routed the rationally selected test, based on residual diagnostics. This avoids relying on a single, potentially poor choice of the test across all proteins. Within this diagnostically curated panel, 13 proteins reached FDR < 0.10, and notably SMOC2, TANK, and IMPG1 were selected by both ML workflows and were among the significant per-protein hits.

Nevertheless, our results remain internal validations to enable us to suggest data-driven hypotheses. External cohorts are required to test whether these convergent proteins retain their discriminative value and to evaluate their incremental contribution to multivariate prediction of CNS side effects due to ASMs.

### Clinical

Our analyses showed that the plasma proteome was indeed different in patients reporting CNS-side effects. In S1 Fig, the pathway and disease enrichments clustered around immune, autoimmune and vascular inflammation pathways. It is possible that CNS-related side effects could be related to such processes; neuroinflammation, BBB dysfunction and microvascular problems. A review in 2017 elaborates on how systemic immune activation disrupts BBB integrity through cytokines and endothelial changes, leading to neuroinflammation and clinical CNS manifestations such as cognitive dysfunction. (17) A study in 2014 showed that activated cytotoxic T cells as part of the adaptive immune responses alone can induce BBB breakdown, neuroinflammation and CNS pathology.(18) Cytokine networks, JAK/STAT signaling, prostaglandins and T-cell responses, all of which were significantly enriched by our protein set, form a mechanistic backbone of both systemic drug hypersensitivity and neuroinflammatory responses in the brain. Cytokine-activated JAK/STAT signaling in microglia and astrocytes, promotes neuroinflammation and synaptic dysfunction besides neurodegeneration in CNS disorders. Apparently, upstream cytokine networks lead to JAK/STAT-driven transcriptional changes in the brain, causing CNS pathology and neuroinflammatory responses (19). A cautious interpretation is that our 61-candidate protein panel may be indicating a pre-existing immune and inflammatory predisposition and vulnerability that changes how the brain and the rest of the body react to ASMs. In such individuals, the normal ASMs effects might be more likely to lead to harsher cytokine responses, BBB perturbation or vascular problems, which increases the likelihood of CNS side effects.

Our results indicate that genes associated with inflammatory processes are commonly differently expressed with CNS-side effects, as several proteins involved in inflammation were significantly expressed among individuals who had CNS-side effects. We found that both TANK and TRIM 40 were differently expressed among individuals with side effects. Both these proteins affect NF-κB (20-22). NF-κB is mainly a pro-inflammatory factor, and as both TANK and TRIM40 and were found significantly increased in our study, more complex pathways than just an increased inflammatory drive may be involved (23).

Furthermore, we found that both IMPG1 and SAG were differently expressed. These proteins are mainly found in the retina and abnormalities are associated with retinal disorders (24, 25). In this study, we included visual side effects as part of the CNS side effects, and the altered protein expression of these proteins are probably more likely found among individuals with visual side effects.

Additionally, we found that NOS1 was differently expressed, a gene that has been associated with a variety of neurologic and psychiatric symptoms, including schizophrenia, neuropsychiatric and neurodegenerative disorders (26). AMFR was also differently expressed and lower levels of AMFR has previously been associated with Alzheimer’s disease.(27) In our cohort, side effects tends to be associated with increased levels; which indicates that also an overexpression of this protein may be associated with neurological symptoms.

A potential limitation is that inflammation is also linked to seizures, which tend to be more frequent in patients with many simultaneous ASMs and more side effects. Further studies are needed to elucidate which of the identified proteins that are specifically associated with CNS-related side effects like dizziness or cognitive symptoms. Larger studies in seizure-free cohorts could be one possibility.

## Materials and Methods

### Study design and participant selection

The Prospective Regional Epilepsy Database and Biobank for Individualized Clinical Treatment (PREDICT) (clinicaltrials.org, NCT04559919) aims to identify biomarkers of clinical utility for detecting treatment effect or side effects in epilepsy and has recruited participants at five neurology outpatient clinics in Västra Götaland region of Sweden since December 2020. The cohort has a similar age and gender distribution to all adult patients seen at neurology departments for epilepsy in the region and Sweden as a whole.(28) The study is approved by the Swedish Ethical Review Authority (approval number 2020-00853). All participants provided informed consent before inclusion as per the Declaration of Helsinki.

### Outcome and variables

Participants in PREDICT fill out a survey with a question on side effects (yes/no) and free text question on the nature of those side effect. Side effects were categorized as CNS-related or not CNS-related by JZ. Information on age and gender was obtained at inclusion.

We analyzed proteomic data from n=176 plasma samples profiled on 1,447 proteins (OLINK Explore384 Neurology I and II and Inflammation I and II panels) in a previous study; after excluding samples with missing information on side effects, n=161 remained. The binary outcome was CNS side-effects (“yes”/“no”). Two covariates, age (numeric) and gender (factor),were used for covariate adjustment.

### Preprocessing

Protein matrices were imputed per-protein using k-nearest neighbors. For the ML workflow, for each protein, we regressed NPX values on age and gender and retained the residuals to remove linear effects of these covariates. Residualized proteins were then standardized to zero mean and unit variance across samples. The outcome was encoded as a factor. Imputation, covariate residualization, and standardization were applied once on the full dataset prior to resampling. Model fitting, hyper-parameter tuning, bootstrap resampling, and label permutations were conducted strictly within resampling splits.

### Least Absolute Shrinkage and Selection Operator (LASSO)

We analyzed the same dataset using LASSO. In the discovery phase, we applied LASSO logistic regression with a 10×10 repeated CV structure. Within each split, we generated B = 30 class-balanced bootstraps of the training set, performed 5-fold inner CV to select the LASSO penalty lambda (AUC-optimized), and recorded whether each protein had a non-zero coefficient. For fixed training data, alpha, and lambda, the glmnet fit is deterministic. However, lambda selection via cv.glmnet depends on the inner-CV fold assignment and random number streams. We therefore treated the full training procedure (including inner CV) as a stochastic resampling step within the stability-selection framework. Within each CV split, if we were to fit LASSO only once per split (no bootstrap), feature j would either be selected split (β_j_ ≠ 0) and (β_j_ = 0)aggregating across 10×10 splits would give a stability resolution of only 1/100. Therefore, we added a class-balanced bootstrap layer inside every discovery CV split.

For each split s, we draw B = 30 class-balanced bootstrap samples and for each bootstrap b, we fit a LASSO with lambda tuned by inner CV and record if protein j is selected. The split-level observed stability for protein j was calculated as:

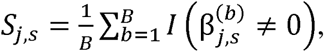

and the observed global stability for protein j is the mean of these proportions across all CV splits:

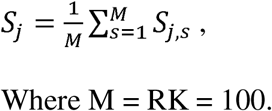

Thus, for each S_j_ we have 3000 resampled fitted models. If the estimated stability for a protein is 0.5 (the minimum acceptable threshold in our study), the Monte Carlo standard error for *S_j_* is:

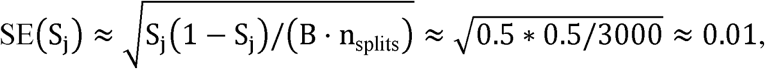

indicating that in compliance with the data-driven goals of this study, stability estimates were numerically precise relative to the decision threshold S_j_ ≥ 0.5. To obtain a null distribution for stability, we repeated the same procedure under B_perm_ = 30 label permutations per split and aggregated permuted selection frequencies. Precisely, within each split s = 1,…, M, for each bootstrap iteration b = 1, …, B_perm_, we used the same class-balanced bootstrap resampling scheme on the split’s training set, but then permuted the bootstrap outcome labels and refitted the LASSO with the same inner 5-fold CV tuning of lambda. (for each split s and permutation index b). Let

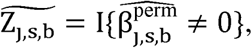

denote the permutation-based selection indicator for protein j in split s and permutation/replicate index b. We then aggregated across splits while keeping the permutation index b intact:

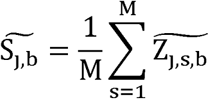

Which is stored in a matrix comprising null stability proportions of dimension (#*proteins*) × B_perm_. In other words, we maintained per-split means and accumulated per-permutation totals across splits and finally divided by M to obtain 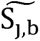. Here, b indexes the within-split bootstrap iteration. For each split s and each b, we generate a class-balanced bootstrap sample (X_boot_, y_boot_), fit an observed LASSO on (X_boot_, y_boot_), and then fit a matched null LASSO on the same X_boot_ but with permuted labels y_perm_ = permute (y_boot_). We then aggregate the permutation indicators across splits separately for each b, yielding B_perm_ null stability statistics 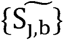 for comparison to the single observed stability S_j_. Then, we computed a Monte Carlo p-value for the hypothesis “protein j is no more stable than expected under random labels” as:

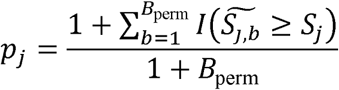

and controlled the false discovery rate (FDR) using the Benjamini–Hochberg (BH) procedure. Proteins with *S_j_* ≥ 0.5 were considered discovery-phase candidates; three proteins met these criteria.

Leak-controlled discovery AUROC for a standard logistic regression and for a LASSO model fitted to all 1447 proteins were obtained by aggregating out-of-fold (OOF) predictions from the 10×10 repeated CV setup and evaluated using pROC. For descriptive purposes, we constructed a permutation-based null for split-level AUROCs by generating permutations, refitting the discovery LASSO across all splits, and plotting the observed and permuted AUROC densities. In an internal validation exploratory phase, we fitted a nested elastic-net logistic model using only the three discovery-phase candidates (train-only median imputation, inner AUC-based tuning of lambda), and reported AUROC from OOF predictions. This nested analysis was explicitly treated as exploratory and not an unbiased estimate of external generalisation.

### Random Forest (RF)

We trained random forest models on a 161 × 1447 residualized proteomic matrix with a binary outcome representing CNS side effects, recoded as *y* ∈ (0, 1) with 1 = “side effects”. Proteins were analyzed in their pre-filtered form; missing values were imputed per split using training-only medians. In the leak-controlled discovery phase we used 10×10 repeated stratified cross-validation (CV) on the full proteomic panel. For each of 100 outer splits, we first tuned RF hyperparameters via an inner CV that maximized Area Under the Receiver Operating Characteristic (AUROC), then trained two models on the outer training set using the tuned hyperparameters: 1) A permutation-importance RF (used only to rank proteins) and, 2) A separate probability RF with a larger number of trees to generate test-set predictions for AUROC estimation.. Within each split, the top 20% of proteins by importance formed a selected set and across all splits, we defined the selection count C_j_ and stability S_j =_ C_j_/100 for each protein j. To assess if these selection stabilities exceeded null expectations, we implemented a symmetric label permutation scenario in which for each permutation *b* = 1,…,*B*_null_ (with *B*_null_ 30), we re-ran the entire discovery phase on permuted labels, recomputed selection counts 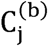, and formed empirical p-values 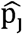 (Monte Carlo estimated p-value (29), not exact p-value) as the proportion of permutations with 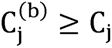:

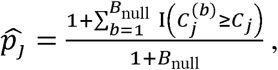

Where I is the indicator function.

In the exploratory phase as the internal validation, we restricted the features to the RF stability-based panel and evaluated its predictive performance with a repeated outer CV with inner hyperparameter tuning on the fixed, pre-defined RF panel (post-selection exploratory internal validation). We tuned RF hyperparameters in the inner loop and obtained subject-level pooled predictions in the outer loop, from which we computed AUROC and DeLong confidence intervals. Split-level AUROC distributions for observed and permuted labels and ROC overlays for discovery versus post-discovery (exploratory) RF were used to visualize model behavior.

### RF and LASSO harmonization

We harmonized LASSO and RF by using the same residualized proteomic matrix, identical stratified 10×10 leak-controlled discovery CV, symmetric label-permutation nulls, and stability-based feature statistics. In both workflows, for each protein, a resampling-based selection frequency, and an empirical p-value were obtained by comparing observed counts to permutation-based null counts, followed by BH FDR. The key difference was how resampling was performed. LASSO is deterministic for a fixed training set, so we engineered an explicit class-balanced bootstrap layer inside each split to obtain a Monte Carlo distribution of selection events and a smooth stability estimate S_j_. RF, however, internally applies bootstrapping and feature subsampling and adding an external bootstrap would double the resampling and would complicate the interpretation. Instead, RF stability was defined as the fraction of outer CV splits in which a protein appears in the top 20% importance fraction. For both LASSO and RF we computed discovery OOF AUROCs using the same repeated CV structure and explicitly fix the ROC direction (positive class = “side efffects”), thereby harmonizing the AUROC interpretation and avoiding artifacts. In both ML workflows given near-random pooled OOF AUROC on the full feature set, permutation-based stability nulls yielded high FDR. Even large numbers of permutation cannot push BH-adjusted FDR well below 0.10. Hence, the choice of data-driven lenient FDR significance thresholds.

### Hierarchical clustering of the data-driven leak-controlled candidate panel proteins

We performed unsupervised hierarchical clustering on the 61 overall candidate proteins identified by both to visualize sample-level proteomic patterns associated with central nervous system (CNS) side effects. Protein expression values were adjusted for age and gender and subsequently standardized (z-scored) across patients. Pairwise Euclidean distances between proteins were computed, and clustering was performed using the complete linkage method. The resulting dendrogram was cut into seven protein clusters, and cluster assignments were visualized as a color-coded top annotation bar. Rows correspond to individual patients. The heatmap was generated using the ComplexHeatmap R package (version 2.24.1), with red representing higher and blue representing lower standardized expression values.

### Per-protein differential expression analysis

We analysed a 61-protein candidate panel (selected upstream by RF stability) to test for group differences between patients with and without CNS side effects. For each protein j for patient i we fitted an ANCOVA model:

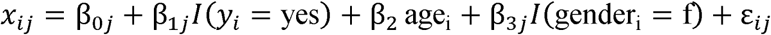

Where I is the indicator function for the side-effect status as the main predictor, x is the NPX values (log2 scale) as the response. We did not scale the NPX values so that we could properly calculate log2 Fold-Change and Fold-Change per protein.

Diagnostic metrics included:

Residual normality: Shapiro–Wilk and Anderson–Darling tests (p ∈ [0,1]); p < 0.05 indicates residuals deviate from Gaussian assumptions.

Variance equality: Brown–Forsythe test (median-based Levene, p ∈ [0,1]); p < 0.05 indicates heteroscedasticity (unequal variance).

Influence: Cook’s distance D_i_ ≥ 0, with D_i_ > 4/(n-p) indicative of influential points; higher values mean greater leverage and impact.

Robust outliers: robust z-scores based on median and MAD (or IQR). |z| > 3 defines extreme values and the “robust outlier fraction” ∈ [0,1] summarizes the proportion of such points.

Based on these diagnostics, each protein was routed to one of three inferential engines:

1. limma moderated t-tests (default) with robust empirical Bayes variance shrinkage, suitable when normality/variance assumptions were reasonable.
2. limma with arrayWeights or HC3 robust SEs when Brown–Forsythe indicated pronounced heteroscedasticity. HC3 inflates standard errors in the presence of unequal variances, yielding conservative p-values.
3. Freedman–Lane permutation ANCOVA (B = 5000) when strong non-normality, outliers or influential points were present and group sizes were modest. Permutation p-values p_perm_∈ [0,1] quantify how extreme the observed group effect is under the null, while preserving the age/gender structure.

Effect sizes were summarised by:

1. Adjusted log2 fold-change (log2FC): the group coefficient, interpretable as the log2 difference between “yes” and “no” at mean age and modal gender. log2FC = 1 implies a 2-fold increase; values in |0.3–0.6| correspond to ∼1.23–1.52× changes.
2. Adjusted fold-change (FC) =2^log2FC^, where FC = 1 implies no change, FC > 1 higher expression in “yes”, FC < 1 lower expression.
3. Hedges’ g, computed on residuals after adjusting for age and gender, with conventional interpretation: |g| < 0.2 negligible, 0.2–0.5 small, 0.5–0.8 moderate, ≥0.8 large; the sign indicates direction of change.

An end-to-end algorithmic description of the three workflows, i.e., LASSO, RF and per-protein differential expression analysis is provided (S1 Supplementary Methods).

### Protein-Protein Interaction (PPI) network analysis

We used the STRING database version 12.0 to subset the PPI network of associations and interactions between the union of the proteins identified by the three workflows (61 proteins). The STRING interactions confidence score cutoff was set to 0.4. We chose the 1^st^ shell of interactors (no more than 10 proteins directly associated with our input proteins) and the 2^nd^ shell of interactors comprising no more than five proteins associated with the 61 input proteins and/or the 1st shell. The mentioned proteins were chosen for the rest of the network and enrichment analyses using STRING online tools. Cytoscape standalone software was used for network metrics analysis and Gephi standalone software was used for visualization and clustering purposes (30, 31). In Gephi, Fruchterman Reingold (32) and Force Atlas 2 (33) layouts were used for visualization purposes. Modularity detection was performed by an algorithm to unfold graph communities using Gephi software suit with randomization to produce a better decomposition (34).

### Software and computation

Analyses were conducted in R version 4.5.0. Parallelization was done using doParallel version 1.0.17, and foreach version 1.5.2. Seeds were fixed for reproducibility.

## Ethical statement

Ethical approval was obtained from the Ethical Review Authority (2020–00853), and the study adhered to the principles outlined in the Declaration of Helsinki. Written informed consent was obtained from all participants before inclusion.

## Acknowledgements

This study was funded by grants from Swedish Research Council, the European Union’s Horizon Europe research and innovation programme under grant agreement No. 101053962, Swedish state through the ALF-agreement (ALFGBG-1006343), ISNF, Margarethahemmet 2025-017 (250601-271231), Swedish Society of Medical Research (SS18-0040), Region Västra Götaland (VGFOUREG-993291), Hjärnfonden FO2025-0024-HK-215, Jeansson foundation, Rune och Ulla Amlövs Stiftelse, Insamlingsstiftelsen för Neurologisk Forskning (ISNF), Anna-Lisa och Bror Björnssons stiftelse, Ann-Louise och Sven-Erik Beiglers stiftelse, Edit Jacobsons Donationsfond 2025-163, Neurofonden, Neuroförbundet and Stiftelsen Ynge Lands minne. JZ reports speaker / advisory board honoraria from UCB, Eisai, Orion Pharma, Angelini Pharma, and Sanofi and as an employee of Sahlgrenska University Hospital being investigator in trials sponsored by SK life science, Bial, UCB, GW Pharma, and Angelini Pharma (no personal compensation).

## Author contributions

S.H.A.: Data curation, Formal analysis, Investigation, Methodology, Software, workflow development, Supervision, Validation, Visualization, Writing original draft

M.K.: Clinical data acquisition and curation, Formal analysis, investigation, Writing original draft

S.A.: Data curation, Experiment design

J.Z.: Conceptualization, Clinical data acquisition and curation, Formal analysis, Funding acquisition, Investigation, Project administration, Resources, Supervision, Writing original draft

## Data availability statement

The proteomics data is protected due to data privacy regulations aimed at protecting sensitive personal information in Sweden. Reasonable requests can be made to the corresponding author and data will be shared if possible, according to a case-by-case evaluation of accordance with Swedish legislation.

## Code reporting

All the codes used to generate the results are in a private GitHub repository. Read access will be granted to the editors and referees upon request.

## Supporting information captions

**S1 Table: LASSO discovery-phase stability-selection statistics for the selected proteins.** For each protein, we report the selection stability, the corresponding mean selection frequency under label permutations (“NullFreq”), the one-sided Monte Carlo p-value p_j obtained from B_perm_ = 30 null permutations (FDR adjusted). Proteins are sorted by decreasing stability and then by increasing FDR

**S2 Table: RF discovery-phase summary of protein-level stability and null-based metrics.** Each row corresponds to a proteomic feature. Columns report the RF selection stability (proportion of discovery splits in which the protein is ranked in the top 20 % of permutation importances), the corresponding selection count across the 100 discovery splits, the empirical Monte Carlo p-value derived from the symmetric label-permutation null adjusted by FDR for all proteins.

**S3 Table: Complete per-protein results for all 61 overall candidate proteins.** The table provides the full output of the data-driven per-protein pipeline for each of the 61 candidate proteins, including the recommended inferential engine, adjusted log□ fold-change, adjusted means in each group, Hedges’ g and FDRs. These results allow readers to evaluate the strength and direction of evidence for all candidates, including proteins that did not reach the predefined FDR threshold but may be of interest for hypothesis generation.

**S4 Table: The PPI network interaction nodes.** STRING PPI edge list and evidence-channel scores for the candidate protein panel. Each row corresponds to an interaction (edge) between node1 and node2, with STRING protein identifiers (node1_string_id, node2_string_id). Columns report per-channel evidence scores from STRING (neighborhood_on_chromosome, gene_fusion, phylogenetic_cooccurrence, homology, coexpression, experimentally_determined_interaction, database_annotated, automated_textmining) and the integrated combined_score representing overall interaction confidence for that edge.

**S5 Table: Gephi community-detection output for the 54-node network.** Nodes (proteins) were visualized in Gephi using the Fruchterman–Reingold and ForceAtlas2 force-directed layouts, and communities (modules) were identified using Gephi’s modularity detection algorithm for unfolding graph communities with randomization to improve the modular decomposition. The algorithm partitioned the network into seven modules (modularity classes). For each node, the table reports its identifier (Id), protein label (Label), occurrence count within the input network/edge list (frequency), node category (type), assigned module (modularity_class), and node eigencentrality (Eigenvector centrality) within the network.

**S1 Fig: Enrichment analyses on all the network proteins.** Gene-Disease, Pathway and Gene Ontology (GO) enrichment results are depicted including the FDR-corrected p-values for each enriched term or pathway. The pathway, Gene Ontology and disease enrichments are mainly clustered around immune, autoimmune and vascular inflammation pathways and ontology terms.

**S1 Supplementary Methods: End-to-end description of the main workflows.** The algorithmic descriptions of the three workflows, i.e., LASSO, RF and per-protein differential expression analysis including the data driven and leak-controlling details.

## Notes

### Competing Interest Statement

Johan Zelano reports speaker / advisory board honoraria from UCB, Eisai, Orion Pharma, Angelini Pharma, and Sanofi and as an employee of Sahlgrenska University Hospital being investigator in trials sponsored by SK life science, Bial, UCB, GW Pharma, and Angelini Pharma (no personal compensation). No other authors report any conflicts of interest.

### Summary of Updates

- The funding and acknowledgment sections have been updated. - Supplementary Figure S1 name has been corrected in the manuscript body.

